# LTK is an ER-resident receptor tyrosine kinase that regulates secretion

**DOI:** 10.1101/575365

**Authors:** Federica G. Centonze, Veronika Reiterer, Karsten Nalbach, Kota Saito, Krzysztof Pawlowski, Christian Behrends, Hesso Farhan

## Abstract

The endoplasmic reticulum (ER) is a key regulator of cellular proteostasis because it controls folding, sorting and degradation of secretory proteins. Much has been learned about how environmentally triggered signaling pathways regulate ER function, but only little is known about local signaling at the ER. The identification of ER-resident signaling molecules will help gain a deeper understanding of the regulation of ER function and thus of proteostasis. Here, we show that leukocyte tyrosine kinase (LTK) is an ER-resident receptor tyrosine kinase. Depletion of LTK as well as its pharmacologic inhibition reduces the number of ER exit sites and slows ER-to-Golgi transport. Furthermore, we show that LTK interacts with and phosphorylates Sec12. Expression of a phosphoablating mutant of Sec12 reduces the efficiency of ER export. Thus, LTK-to-Sec12 signaling represents the first example of an ER-resident signaling module the potential to regulate proteostasis.

## Introduction

The secretory pathway handles a third of the proteome (Sharpe et al., 2010) and it is becoming increasingly clear that its functional organization is regulated by a wide range of signaling pathways (Farhan and Rabouille, 2011; Farhan et al., 2010; Scharaw et al., 2016; Zacharogianni et al., 2011) (Cancino and Luini, 2013; Pulvirenti et al., 2008). Much has already been learned about how the secretory pathway responds to external stimuli. However, our understanding of its autoregulation, i.e. about its response to stimuli from within the endomembrane system, is less developed. This is mainly due to absence of many examples of signaling cascades operating locally on the secretory pathway. The probably best-understood example for autoregulation of the secretory pathway is the unfolded protein response (UPR). The UPR is induced by an accumulation of unfolded proteins in the endoplasmic reticulum (ER), which results in increasing the expression of chaperones as well as the machinery for protein degradation, vesicle budding, tethering and fusion (Gardner et al., 2013). Another example for local signaling at the secretory pathway is activation of Src family kinases Gαq at the Golgi and the ER (Giannotta et al., 2012; Pulvirenti et al., 2008; Subramanian et al., 2019). As for Src family kinases, a large pool of them localizes to the Golgi. However, as far as the ER is concerned, no resident signaling molecules are known to permanently or dominantly localize to this cellular location, apart from the classical UPR signal transducers. In fact, the only known examples of ER-localized signaling molecules are limited to mutant or oncogenic variants of signaling molecules (Choudhary et al., 2009; Schmidt-Arras et al., 2009). Thus, signaling at the ER remain poorly understood, which emphasizes the importance of the quest for ER-localized or -resident signaling molecules.

COPII vesicles that form at ER exit sites (ERES) are responsible for ferrying secretory cargo out of the ER. The COPII coat is composed of the small GTPase Sar1, the Sec23-Sec24 heterodimer and the Sec13-Sec31 heterotetramer (Zanetti et al., 2011). Activation of Sar1 is mediated by its exchange factor, Sec12, a type-II transmembrane protein that localizes to the general ER as well as to ERES(Montegna et al., 2012; Saito et al., 2014). ERES were discovered as COPII decorated sites that often localize in close vicinity to the ER Golgi intermediate compartment (ERGIC) (Appenzeller-Herzog and Hauri, 2006; Orci et al., 1991).

Previous siRNA screens uncovered of a collection of kinases that regulate ERES (Farhan et al., 2010; Simpson et al., 2012). Here, we focused on leukocyte tyrosine kinase (LTK) because it was the only hit in our previous screen that was reported to partially localize to the ER (Farhan et al., 2010) (Bauskin et al., 1991). Our current work identifies LTK as the first ER resident receptor tyrosine kinase that regulates COPII dependent trafficking and is thus a potential druggable proteostasis regulator.

## Results

### LTK localizes to the ER

LTK is a receptor tyrosine kinase that is highly homologous to the anaplastic lymphoma kinase (ALK) (Figure 1A). While their cytoplasmic kinase domain is 79% identical, the extracellular domain of ALK is much larger than that of mammalian LTK as it contains two MAM domains (acronym derived from meprin, A-5 protein, and receptor protein-tyrosine phosphatase mu). Analysis of LTK and ALK evolution shows that deletions of the largest part of the extracellular domain of LTK occurred only in mammals (Figure 1B). Non-mammalian LTK rather resembles ALK than human LTK.

**Figure 1.**
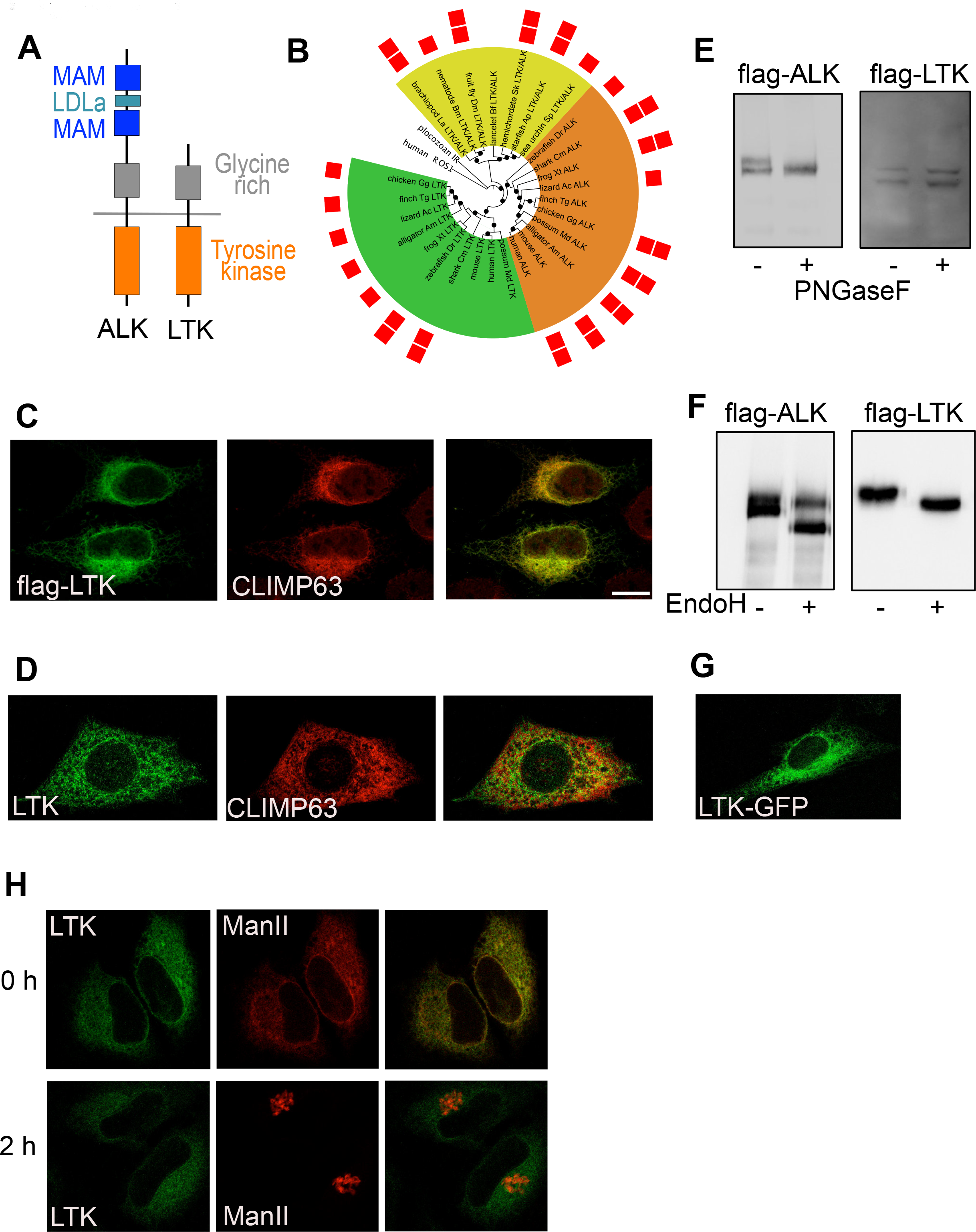
Subcellular localization of LTK. ***A***, Schematic illustrating the domains of LTK and ALK. ***B***, Phylogenetic tree of ALK and LTK kinase domains. Color ranges highlight invertebrate LTK/ALK-like proteins (yellow), vertebrate ALK proteins (orange) and vertebrate LTK proteins (green). Red squares indicate presence of one or two MAM domains. Black circles mark branches with bootstrap support above 50%. Human ROS1 and placozoan insulin receptor-like kinase domains used as outgroup. The list of abbreviations used in the figure are found in the supplementary dataset. ***C***, Immunostaining of flag-tagged human LTK and endogenous CLIMP63 in HeLa cells. ***D***, immunofluorescence staining of endogenous LTK and CLIMP63 in HepG2 cells. ***E***, HeLa cells expressing flag-tagged LTK or ALK were treated with PNGaseF followed by lysis and immunoblotting against flag to detect ALK or LTK. ***F***, HeLa cells expressing flag-tagged LTK or ALK were lysed and the lysate treated with endoH followed by immunoblotting against flag to detect ALK or LTK. ***G***, HeLa cells expressing GFP-tagged LTK were imaged using live microscopy. ***H***, HeLa cells expressing GFP-tagged LTK and mCherry tagged ManII in the RUSH system were treated for 0 or 2 h with biotin followed by fixation.

LTK was reported to localize to the ER (Bauskin et al., 1991), but this was questioned by recent findings showing LTK activation by extracellular ligands (Reshetnyak et al., 2015; Zhang et al., 2014). Overexpressed flag-tagged LTK, but not ALK colocalized with the ER marker CLIMP63 (Figure 1C, Figure S1A). Endogenous LTK also localized to the ER (Figure 1D). The specificity of the antibody was tested by showing that the fluorescence signal is weaker in LTK depleted cells (Figure S1B). We also noticed in 10% of cells a weak colocalization of LTK with the ERES marker Sec31 (Figure S1C). Endogenous LTK was analyzed by immunofluorescence was carried out in HepG2, because they express high levels of LTK, but are essentially ALK negative (Figure S2A), which limits the possibility of antibody crosstalk.

To corroborate the immunofluorescence results, we subjected intact cells expressing flag-tagged ALK or LTK to PNGaseF, an enzyme that cleaves glycans and is therefore expected to cause a shift in electrophoretic mobility. Consistent with its absence at the cell surface, we found that LTK was insensitive to treatment of cells with PNGaseF (Figure 1E). On the contrary, ALK was sensitive to digestion with PNGaseF (Figure 1E), which is consistent with its expression at the cell surface. We next treated cell lysates expressing flag-tagged LTK or ALK with endoglycosidase H (EndoH), which only digests core-glycosylated proteins that have not entered the Golgi apparatus. LTK was completely sensitive to EndoH treatment, indicating that it resides in a pre-Golgi compartment (Figure 1F). On the other hand, only 60% of the ALK pool was sensitive to EndoH (Figure 1F). Available antibodies do not detect endogenous LTK by immunoblotting, preventing us from performing the same analysis with endogenous LTK.

To rule out that the absence of staining of LTK at the cell surface is due to fixation artifacts, we tagged LTK with GFP and performed live imaging. LTK localization was similar as in fixed cells, and was reminiscent of the ER (Figure 1G). Finally, we wanted to directly test whether LTK leaves the ER using the retention using selective hooks (RUSH) assay (Boncompain et al., 2012). The RUSH assay monitors the trafficking of a fluorescently-labeled reporter protein out of the ER. This reporter is retained in the ER through a streptavidin-based interaction with an ER-resident hook. Treatment with biotin relieves retention and allows the reporter to exit towards post-ER compartments. We engineered GFP-tagged LTK into the RUSH system and expressed it together with a well-described secretory RUSH reporter, Mannosidase-II (Man-II) tagged with mCherry. Initially, LTK and Man-II colocalized in the ER (Fig. 1H). Cells were followed for 2 h after biotin addition, a time point at which Man-II was entirely in the Golgi. However, LTK was still ER-localized (Figure 1H), indicating that it does also not leave the ER under synchronized trafficking conditions. Altogether, our results show that LTK is an ER-resident receptor tyrosine kinase, making it a promising candidate to regulate secretion by local ER-based signaling.

### LTK regulates ER export

The ER localization of LTK prompted us to ask whether it might regulate ER-to-Golgi trafficking. Knockdown of LTK (Figure S2B) resulted in a reduction of the number of ERES by 30-40% in HepG2 (Figure 2A&B) and HeLa cells (Figure S2C). To support the results of the knockdown experiments, we treated HepG2 cells with two LTK inhibitors, alectinib and crizotinib. Of note, HepG2 and HeLa cells are essentially ALK negative (Figure S2A), and thus any effect of these drugs is due to LTK inhibition. Treatment with both drugs for 30 minutes resulted in a reduction in the number of ERES comparable to LTK knockdown (Figure 2B). Notably, crizotinib had no effect on ERES in in LTK depleted cells, supporting the notion that it acts via LTK inhibition (Figure 2C). We confirmed that crizotinib and alectinib inhibited LTK autophosphorylation in our experimental system (Figure 2D). We also tested the effect on crizotinib in live imaging and found that the onset of ERES reduction is after approximately 10 min of treatment (Figure 2E). To determine the effect on ER-to-Golgi trafficking, we used the RUSH assay with mannosidase-II as a RUSH cargo (RUSH-ManII) (Boncompain et al., 2012). Silencing LTK expression or its chemical inhibition resulted in a clear retardation of trafficking to the Golgi (Figure 3A, Supplementary movies S1&2). This effect was rather a retardation of traffic, than a total inhibition, because when we allowed the RUSH cargo to traffic for two hours, there was no difference between control and LTK inhibited cells (Figure 3B). The effect of LTK knockdown was not limited to Man-II, but was also observed with another RUSH cargo, namely collagen X (Figure 3C), indicating that the effect of LTK is not limited to one type of cargo. Our results so far indicate that LTK is an ER-resident receptor tyrosine kinase that regulates ER export.

**Figure 2.**
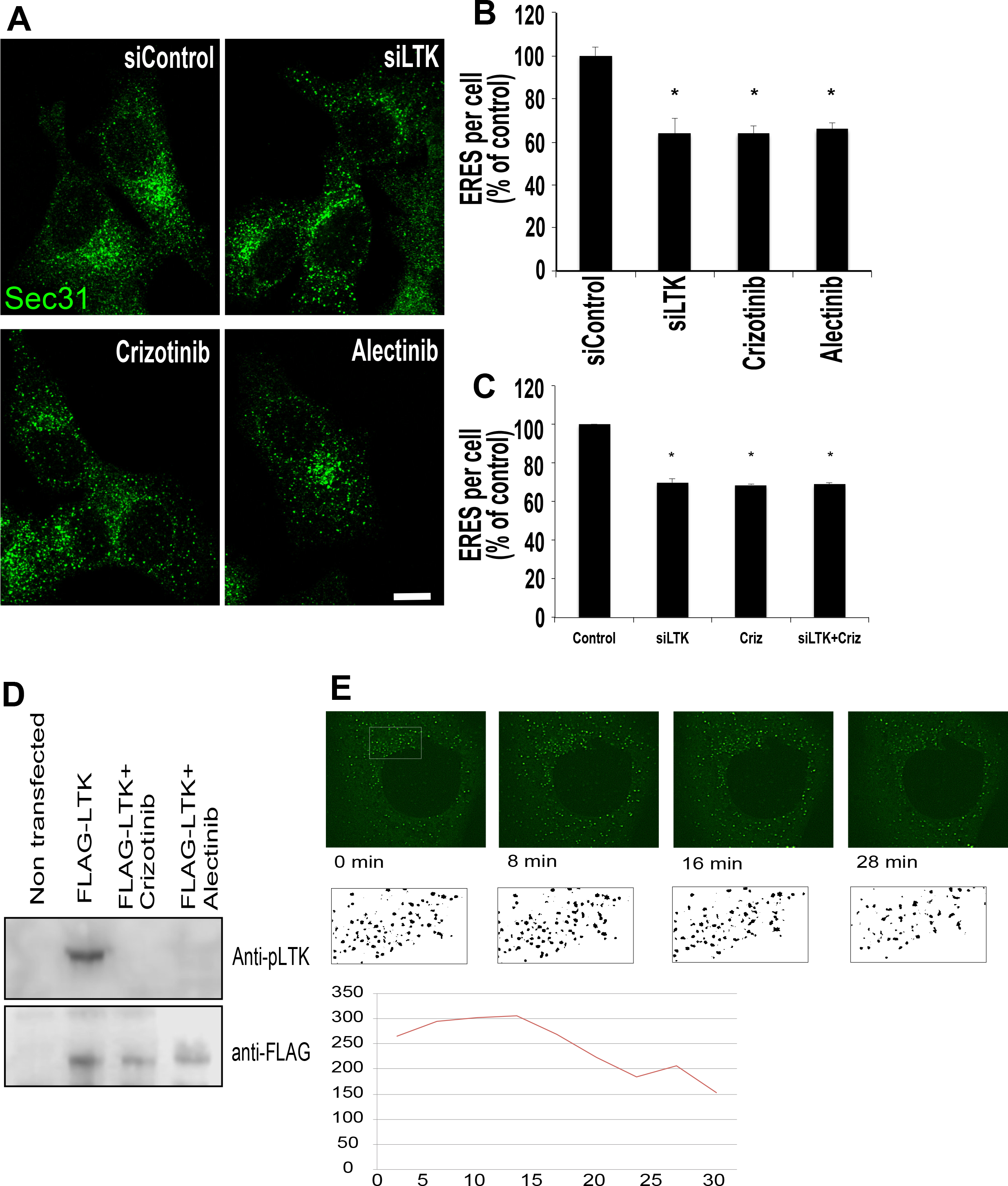
LTK regulates ER export and ERES. ***A**,* HepG2 cells were either subjected to LTK silencing with siRNA followed by fixation and staining after 72 h. Alternatively, cells were treated with crizotinib and alectinib (1μM) for 30 min prior to fixation and immunostaining against Sec31 to label ERES. ***B***, Bar graphs on the right side of the panel indicate the efficiency of LTK knockdown (upper graph) and the quantification of ERES number per cell displayed as % of control (control siRNA for the LTK knockdown or solvent treatment for crizotinib and alectinib). Data are from 3 independent experiments with at least 35 cells per experiment per condition. Asterisk indicates statistically significant differences at p<0.05. ***C***, HepG2 cells were transfected with control or LTK siRNA. After 72 h, cells were treated with solvent or with crizotinib (1µM) for 30 min prior to staining for Sec31 to determine ERES number. ***D***, HeLa cells expressing flag-tagged LTK were treated with solvent or with crizotinib or alectinib for 30 min prior to lysis and immunoblotting as indicated. P-LTK indicates immunoblotting against an antibody that detect phosphorylation on Y672. Flag immunoblotting was performed to determine equal loading. ***E***, HeLa cells expressing GFP-Sec16A treated with 1µM crizotinib followed by confocal live imaging. Stills of the indicated time points are depicted. The ERES in the boxed area are depicted in B/W to enhance visibility. The number of ERES was counted and is displayed in the lower graph.

**Figure 3.**
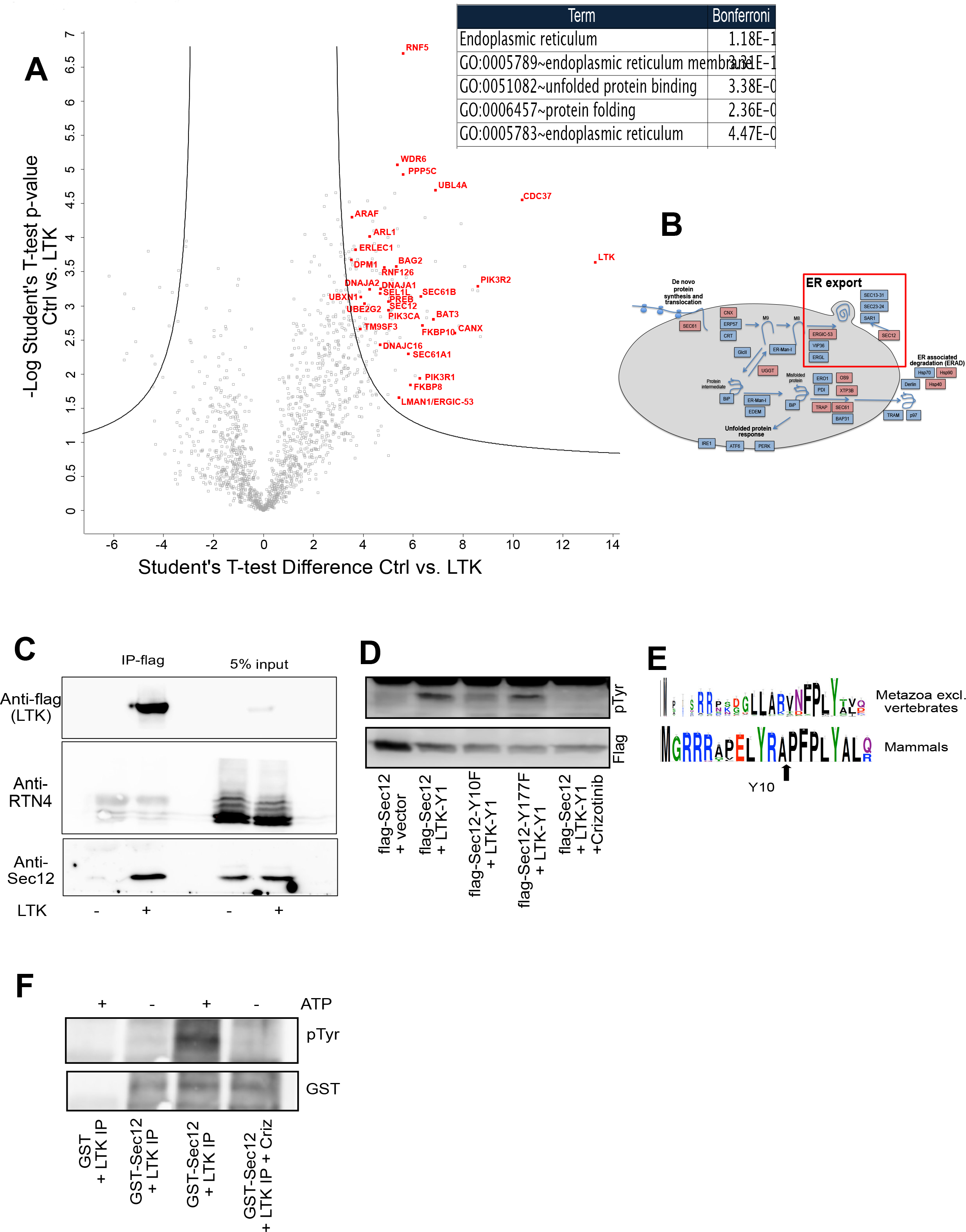
LTK regulates ER export. ***A***, Representative images of HeLa cells stably expressing the GFP-RUSH-ManII construct (Str-KDEL-ManII-EGFP) under different conditions: 0 min, cells not treated with biotin; 20 min, cells fixed 20 minutes after biotin treatment; Cont, control siRNA tranfected; siLTK, LTK silenced; crizotinib and alectinib indicate cells treated with 1μM 30 min prior to biotin addition. Bar graph shows quantification from 3 independent experiments. Asterisk indicates statistically significant differences at p<0.001. Scale bars in this figure are 15 μm. ***B***, Representative images of HeLa cells stably expressing the GFP-RUSH-ManII construct (Str-KDEL-ManII-EGFP) imaged 2 h after release of the reporter from the ER. Two conditions are depicted, control and LTK knockdowen cells. ***C***, HeLa cells expressing the RUSH-Collagen-X construct were transfected with control or LTK siRNA. After 72 h, cells were treated with biotin and fixed immediately (T0) or after 20 min. Cells were immunostained against Giantin to label the Golgi. The increase in green fluorescence in the Golgi region relative to outside the Golgi region was measured using ImageJ. The bar graph represents the mean of 4 independent experiments.

### LTK interacts with and phosphorylates Sec12

We next sough to mechanistically uncover how LTK regulates ER export. To this end, we mapped the interactome of flag-tagged LTK expressed in HEK293 cells, because they do not express endogenous LTK and can be transfected easily. The interactome of LTK uncovered proteins such as known proteins downstream of receptor tyrosine kinases, but also most notably, early secretory pathway proteins such as quality control or trafficking machinery (Figure 4A&B, Table S1). This is consistent with the localization of LTK to the ER. The top associated GO-term among the LTK interactome was “Endoplasmic reticulum” (Figure 4A). Among the potential LTK interaction partners identified, we focused on Sec12 (also known as PREB), due to its well-known role in ER export and the biogenesis of ERES (Barlowe and Schekman, 1993; Montegna et al., 2012; Saito et al., 2014). Sec12 is a type-II transmembrane protein that acts as GEF for Sar1. Using co-immunoprecipitation, we confirmed that LTK interacts with Sec12, but not with an unrelated transmembrane protein of the ER that was not recovered in the interactome (Figures 4C & S3A). Co-expression with LTK resulted in an increase in the tyrosine phosphorylation of Sec12, which was crizotinib sensitive (Figure 4D). Two tyrosine residues in Sec12 (Y177 and Y10) were predicted by databases to be phosphorylated (PhosphoSitePlus and NetPhos3.1). Therefore, we mutated both tyrosine residues to phenylalanine. Mutation of Y10 in Sec12 strongly reduced tyrosine phosphorylation, indicating that this is a strong candidate site for phosphorylation by LTK (Figure 4D). Mutation of Tyrosine 177 had no effect. Tyrosine 10 in Sec12 is a site conserved in mammals but not in non-vertebrates (Figure 4E). We next purified the cytosolic domain of Sec12 and incubated it with flag-LTK immunoprecipitate. Addition of ATP to the mix resulted in a crizotinib-sensitive increase of Sec12 phosphorylation (Figure 4F), supporting the notion that Sec12 phosphorylation is LTK-dependent. We also tested the possibility that tyrosine phosphorylation is dependent on Src family kinases, and found that their inhibition only marginally affected Sec12 tyrosine-phosphorylation (Figure S3B).

**Figure 4.**
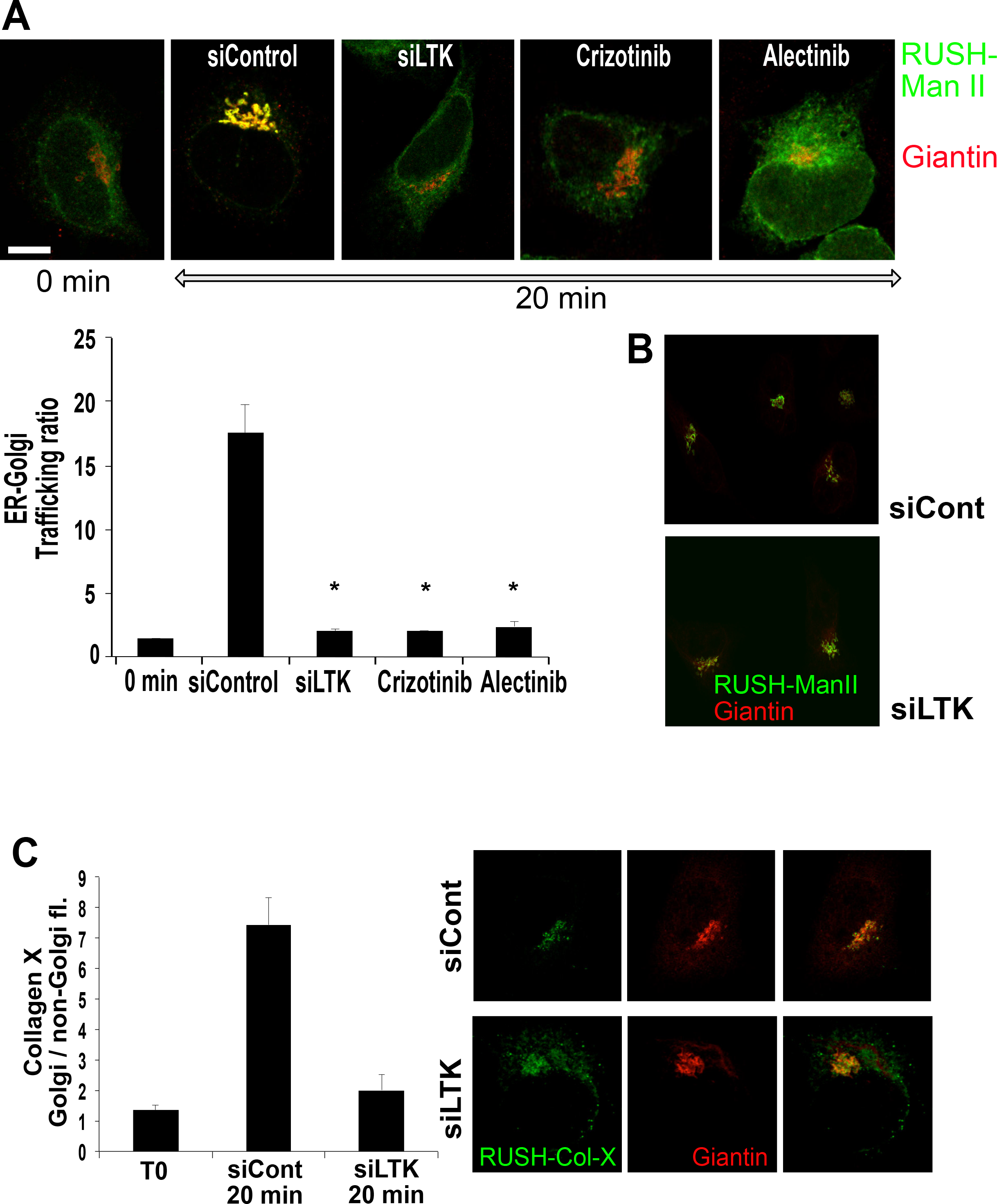
LTK interacts with and phosphorylates Sec12. ***A**,* Volcano-plot of the interactome of HA-tagged LTK revealed by IP-MS from HEK293 cells. The red labeled candidate interacting proteins are related to RTK signaling or ER-associated processes. The table indicates the top-scoring biological processes enriched among the LTK interaction partners. ***B***, Schematic representation of key components of the ER folding, quality control and export machinery with red-highlighted interaction partners. Red box highlights ER-export. ***C***, Flag-tagged LTK was immunoprecipiated followed by immunoblotting against Sec12, which was identified in the interactome and against RTN4, a transmembrane protein that we did not find in the LTK interactome. ***D**,* HeLa cells were transfected with vectors encoding flag-tagged Sec12 or its mutants together with an empty vector or with YFP1-tagged LTK (LTK-Y1). In the last lane are lysates from cells pre-treated with 1μM crizotinib for 30 min. ***E***, Sequence logo to demonstrate the conservation of amino acids in Sec12 in mammals or in metazoan excluding vertebrates. ***F***, Purified GST-tagged cytosolic domain of Sec12 or GST were incubated a flag-LTK immunoprecipitate from HEK293 cells (LTK IP) in the presence or absence of ATP or crizotinib.

Because Sec12 is the exchange factor for Sar1, we next tested the effect of LTK inhibition on the dynamics of YFP-tagged Sar1A using fluorescence recovery after photobleaching (FRAP) microscopy. Cells were pretreated with solvent or with crizotinib for 20 minutes prior to FRAP microscopy. Inhibition of LTK reduced the mobile fraction of Sar1, indicative of a reduced exchange activity on single ERES (Figure 5D). No effect of crizotinib on general ER structure was detected (Figure S3C). We tried using the tryptophan fluorescence assay to monitor GTP exchange in Sar1 and its modulation by LTK. However, this assay cannot be used because the inclusion of ATP to the reaction (to promote Sec12 phosphorylation) distorted the assay (data not shown). Thus, we used a different approach to support the results of the FRAP assay, namely by immunostaining for Sar1-GTP positive ERES. This approach has been validated by other previously (Venditti et al., 2012). Treatment of cells with crizotinib resulted in cells with less and fainter Sar1-GTP positive puncta (Figure 5B), indicating that inhibition of LTK affects the recruitment of Sar1 to ERES.

**Figure 5.**
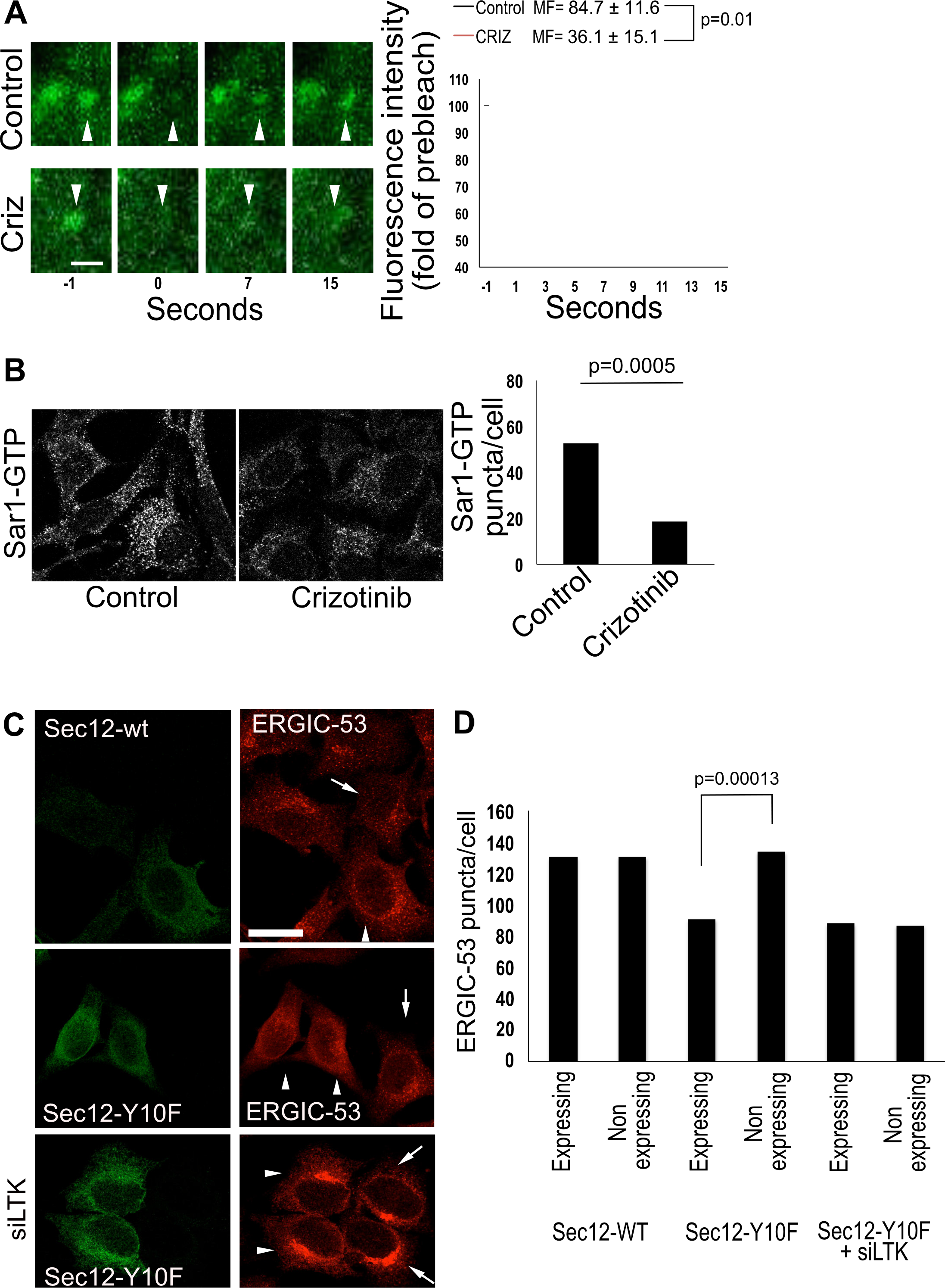
LTK regulates Sec12 function. ***A**,* FRAP assay of HepG2 cells expressing YFP-tagged Sar1A. Images on the left side show magnified single ERES at different time points before (−1) and directly after (0) bleaching as well as at the indicated time points after bleaching. Graph shows an evaluation of 9 FRAP curves for each condition from three experiments. MF= mobile fraction. Scale bar= 1 μm. ***B**,* HeLa cells were treated with solvent (Control) or 1μM crizotinib for 30 min prior to fixation and immunostaining against Sar1-GTP. The number of Sar1-GTP puncta was counted using ImageJ and is displayed in the bar graph on the right side of the panel. Results represent the average number of puncta per cell obtained from 100-150 cells. Statistical significance was tested using unpaired, two-tailed t-test. ***C***, Wild type Sec12 or its mutant Sec12-Y10F were expressed in HeLa cells immunostained for ERGIC-53 and flag. siLTK= expression of Sec12 in LTK depleted cells. Arrows indicate non-transfected cells. Arrowheads indicate cells expressing Sec12. ***D**,* Bar graph showing the quantification of the number of ERGIC-53 puncta per cell from three independent experiments. The graph compares cells expressing the Sec12 construct to directly adjacent non-transfected cells. Scale bar= 30 μm.

To obtain further support for a role of Sec12 phosphorylation in ERES function, we determined the number of peripheral ERGIC-53 structures in cells expressing a phosphoablating mutant of Sec12 (Sec12-Y10F). Peripheral ERGIC structures are good indicators of ERES function (Ben-Tekaya et al., 2005; Farhan et al., 2010). Expression of Sec12-Y10F resulted in a decrease in the number of ERGIC-53 punctae (Figure 5C). No effect of Sec12-Y10F expression was detected in LTK knockdown cells (Figure 5C). Altogether, we propose that Sec12 is phosphorylated in a manner dependent on LTK and that this phosphorylation affects ERES function.

Over the past decade, mounting evidence indicates that endomembranes are subject to regulation by signaling pathways (Baschieri et al., 2014; Cancino and Luini, 2013; Farhan and Rabouille, 2011). An emerging concept of endomembrane signaling is autoregulation, which is defined as a response of a biologic system that help re-establish homeostasis. The UPR is the best understood and characterized autoregulatory response of the secretory pathway (Gardner et al., 2013; Ron and Walter, 2007) making it a useful template to compare other autoregulatory circuits with. The UPR is induced by misfolded or unfolded proteins and its main purpose is to globally upregulate the capacity of the endomembrane system to promote protein folding or their degradation. Such a broad response is expected because the purpose of the UPR is to maintain or re-establish global homeostasis of the ER. Another feature of the UPR is that its main sensors and mediators such as IRE1 or ATF6 are resident to the ER. Contrary to the response of the ER to misfolded proteins, we know very little whether and how local signaling at the ER controls the capacity of the ER to unload of folded proteins, *i.e.* of ER export. Very recently, a signaling cascade including Gαq was shown to operate at ERES (Subramanian et al., 2019). However, this signaling circuit control the export of a small subset of proteins and interfering with it had no effect on general protein secretion or on global ERES number. Thus, this novel pathway represents a tailored response of the ER, which is unlike the more global response of the UPR. Another difference to the UPR is that the main signaling mediator of this pathway Gαq is not resident at the ER or ERES.

As far as LTK is concerned, our results indicate that it might be more similar to the UPR. Firstly, LTK is resident in the ER and secondly, the effect of LTK is a global regulation of trafficking and ERES number. This is in line with the observation that LTK phosphorylates Sec12, a general regulator of ERES biogenesis. Future work will need to address the question concerning what stimuli activate LTK. Because the LTK interactome contained several cargo receptors such as ERGIC-53, VIP36, ERGL, ERGIC1 and SURF4, we speculate that these cargo receptors might represent stimuli that induce LTK activity to positively regulate ER export. In fact, our preliminary data indicate that clients of ERGIC-53 positively regulate LTK activity (Centonze and Farhan, nonpublished data).

The role of LTK in secretion might also be relevant for human diseases. Gain of function mutations in LTK have been observed in patients and mice with systemic lupus erythematosous (Li et al., 2004). We speculate that this gain of function mutation confers a selective advantage to autoimmune plasma cells as it allows them to cope with a higher secretory load. LTK might also represent a suitable drug target in cancer therapy, especially since cancer cells are considered to be addicted to secretion due to a high proteostatic challenge (Dejeans et al., 2015; Urra et al., 2016). This notion is supported by our observation that LTK inhibition increases the ER stress response (as measured by increased XBP1s levels) in cells treated with thapsigargin (Figure S3D). The investigation of the potential of LTK as a drug target will be an interesting area of future investigation.

## Materials & Methods

### Mass spectrometry

Immunoprecipitated proteins were eluted from beads using repeated incubations with 8 M Guanidinhydrochlorid (GdnHCl) at pH 8 and subjected to reductive alkylation (using 15 mM iodoacetamide and 5 mM DTT) and methanol/chloroform extraction followed by digestion with sequencing-grade Trypsin (Promega) overnight at 37°C.Tryptic peptides were desalted and analyzed by liquid chromatography tandem mass spectrometry (LC-MS/MS) using a NanoLC 1200 coupled via a nano-electrospray ionization source to a Q Exactive HF mass spectrometer. Peptide separation was carried out according to their hydrophobicity on an in-house packed 18 cm column with 3 mm C18 beads (Dr. Maisch GmbH) using a binary buffer system consisting of solution A (0.1% formic acid) and B (80% acetonitrile, 0.1% formic acid). Linear gradients from 7 to 38% B in 35 min were applied with a following increase to 95% B within 5 min and a re-equilibration to 5% B. MS spectra were acquired using 3e6 as an AGC target, a maximal injection time of 20 ms and a 60,000 resolution at 200 m/z. The mass spectrometer operated in a data dependent Top15 mode with subsequent acquisition of higher-energy collisional dissociation fragmentation MS/MS spectra of the top 15 most intense peaks. Resolution for MS/MS spectra was set to 30,000 at 200 m/z, AGC target to 1e5, max. injection time to 64 ms and the isolation window to 1.6 Th. Raw data files were processed with MaxQuant (1.6.0.1) as described previously (Cox and Mann, 2008; Cox et al., 2011) using human (UP000005640) UNIPROT databases, tryptic specifications and default settings for mass tolerances for MS and MS/MS spectra. Carbamidomethylation at cysteine residues was set as a fixed modification, while oxidations at methionine and acetylation at the N-terminus were defined as variable modifications. The minimal peptide length was set to seven amino acids and the false discovery rate for proteins and peptide-spectrum matches to 1%. Perseus (1.5.8.5) was used for further analysis (Pearson’s correlation, two-sample t-test) and data visualization (Tyanova et al. 2016). Functional annotation enrichment analysis was performed using DAVID (Huang et al., 2007) coupled to significance determination using Fisher’s exact test and correction for multiple hypothesis testing by the Benjamini and Hochberg FDR.

### Immunofluorescence

Cells were fixed in 3% paraformaldehyde for 20 min at room temperature. Afterwards, cells were washed in PBS with 20 mM glycine followed by incubation in permeabilization buffer (PBS with 0.2% triton X100) for 5 min at room temperature. Subsequently cell were incubated for 1 h with the primary antibody and after washing for another 1 h in secondary antibody diluted in 3% BSA in PBS. Cells were mounted in polyvinylalcohole with DABCO antifade and imaged.

For Sar1-GTP staining, cells were fixed and permeabilized in ice-cold 50% methanol-50% acetone for 10 minutes at −20. Subsequently cells were incubated for 1 h with the blocking buffer (PBS with 10% goat serum) at room temperature followed by incubation with primary antibody for 2 hours and another 1 hour with secondary antibody, both diluted in 3% BSA in PBS.

### Cell culture and transfection

HeLa, HEK293T and HepG2 cells were cultured in DMEM (GIBCO) supplemented with 10% FCS and 1% Penicillin/Streptomycin (GIBCO).

For overexpression of plasmids, cells were transfected with either Fugene 6 or with TransIT-LT1 (Mirus). For knockdown experiments, cells were reverse transfected with 10nM siRNA (final concentration) using HiPerfect (Qiagen) according to manufacturer’s instruction.

### Cell lysis, immunoblotting and immunoprecipitation

Cells were washed twice with PBS and collected in lysis buffer (50mM Tris-HCl, pH7.4; 1mM EDTA, 100mM NaCl, 0.1% SDS and 1% NP40) supplemented with proteinase and phosphatase inhibitor (Pierce Protease and Phosphatase Inhibitor Mini Tablets, EDTA-free). Lysates were incubated on ice for 10minutes followed by clearing centrifugation at 20000xg at 4°C for 10 min. Supernatants were transferred into fresh tube and reducing loading buffer was added. Lysates were subjected to SDS page and transferred on a nitrocellulose membrane using semidry transfer. The membrane was blocked (in ROTI buffer (Roth) or 5% milk in PBS with 0.1% tween) and probed with the appropriate primary antibodies. Subsequently membranes were incubated with horseradish peroxidase conjugated secondary antibody. Immunoblots were developed using a chemiluminescence reagent (ECL clarity, BioRad) and imaged using ChemiDoc (BioRad).

For immunoprecipitation experiments, cells were lysed in IP-buffer (20mM Tris-HCl, pH 7.4, 150mM NaCl, 1mM MgCl2, 10% glycerol, 0.5% NP40, n-Dodecyl-B-D-maltoside).

### PGNase F digestion and EndoH

For PGNase F digestion, 3.2 x10^5^ HeLa cells were seeded into 6-well plates and the next day transfected with 1 μg plasmid DNA. 24 h later, cells were washed with PBS and the cells were incubated in 1ml serum free medium and 250U/ml PNGase F (NEB, P0704S) for 6h. Subsequently, cells were lysed as described above.

For EndoH digestion, 2×10^6^ cells were plated into in a 10 cm dish. After 24 h, cells were transfected with 3ug plasmid DNA. The next day, cells were lysed in IP buffer (50mM Tris-HCl pH 7.4, 10% glycerol, 150mM NaCl, 2mM EDTA, 0.5% Tx-100) supplemented with proteinase and phosphatase inhibitor (Pierce Protease and Phosphatase Inhibitor Mini Tablets, EDTA-free). Immunoprecipitation against flag was performed using EZview™ Red ANTI-FLAG^®^ M2 Affinity Gel (Sigma) overnight at 4°C followed by washing in IP buffer. Beads were incubated with 1000 units EndoH (P0702S, NEB) according to manufacturer’s instructions for 90 min at 37°C. Subsequently, the reaction was stopped by adding reducing sample buffer.

### Imaging

Imaging was performed on laser scanning confocal microscopes: LeicaSP5 and Zeiss LSM700. All images were acquired using a 63x oil immersion objective (NA 1.4).

FRAP was performed on a LeicaSP5 confocal microscope using a 63×/1.4 NA oil-immersion objective at 3-fold digital magnification. All experiments were performed at 37°C and cell were maintained in complete medium supplemented with 25mM HEPE (pH7.4). After acquisition of a pre-bleach image, the ERES was bleached at 100% laser intensity for 750 ms. After bleaching, images were acquired at one image per second. Images were analyzed using ImageJ. The fluorescence intensity of the ERES prior to bleaching was set to 100% and all subsequent values were normalized to it. The mobile fraction was calculated as MF=(F_∞_−F_0_)/(F_i_−F_0_), where F∞ is fluorescence in the bleached region after recovery, Fi is the fluorescence in the bleached region before bleaching, and F0 is the fluorescence in the bleached region directly after bleaching.

ERES were quantified as described previously (Tillmann et al, 2015).

### Sequence analysis

The sequences of ALK and LTK kinase domains were aligned using Muscle algorithm and Jalview environment (Edgar, 2004; Waterhouse et al. 2009). Phylogenetic trees were built using the PhyML program (Guindon et al. 2009) on the phylogeny.fr server (Dereeper et al. 2008) and visualised with the help of the iToL server (Letunic and Bork, 2016). Protein domains were detected using CD search (Marchler-Bauer et al. 2017). The following abbreviations were used in Figure 1B: alligator Am, Alligator mississippiensis; brachiopod La, Lingula anatine; chicken Gg, Gallus gallus; finch Tg, Taeniopygia guttata; frog Xt, Xenopus tropicalis; fruit fly Dm, Drosophila melanogaster hemichordate Sk, Saccoglossus kowalevskii; lancelet Bf, Branchiostoma floridae; lizard Ac, Anolis carolinensis; mouse Mm, Mus musculus; nematode Bm, Brugia malayi; plocozoan Ta, Trichoplax adhaerens; possum Md, Monodelphis domestica; sea urchin Sp, Strongylocentrotus purpuratus; shark; Cm, Callorhinchus milii; starfish Ap, Acanthaster planci; zebrafish Dr, Danio rerio

### Purification of GST-tagged Sec12 cytosolic domain

E.Coli BL21 (DE3) were transformed with GST-tagged Sec12 construct. Bacteria were cultured in HSG growth medium and induction was performed with 0.4 mM IPTG. Bacterial pellet was dissolved in lysis buffer (50mM Tris pH8, 150 NaCl, 10% glycerol, 0.1% Triton-X100, 100ug/ml lysozyme supplemented with proteinase and phosphatase inhibitor), sonified and centrifuged at 32000xg for 30 minutes at 8°C. Supernatant was incubated with Gluthathione Sepharose 4 Fast Flow (GE Healthcare) for 2 hours at 4 °C, washed with PBS and beads were resuspended in buffer (50 mM Tris-HCl pH 7.5, 150 Mm NaCl, 5% glycerol).

### Kinase assay

HeLa cells expressing flag-tagged LTK were lysed and LTK was immunoprecipitated using anti-flag M2 beads. The immunoprecipitate was re-suspended in buffer (20mM Tris-HCL, pH7.4, 150 mM NaCl, 10mM MgCl_2_, 10% glycerol). Typically, 5.10^6^ cells were used. The immunoprecipitate was incubated with 1.5 μg of GST or GST-tagged Sec12 cytosolic domain for 30 at 30°C. To induce kinase activity 400 μM ATP was included. The reaction was stopped by adding sample buffer.

### qPCR

The levels of LTK and ALK were determined by qRT-PCR. Total RNA was extracted using Direct-Zol RNA kit (Zymo Research, Irvine, CA, USA) and cDNA was reverse-transcribed using High Capacity cDNA Reverse Transcription Kit (Thermo Fisher Scientific, Waltham, MA, USA). The expression of LTK and ALK was determined using LightCycler 480 SYBR Green I Master Mix (Roche Life Science, Switzerland) and normalized to GAPDH using commercially available primers (Qiagen, Germany; for LTK: QT00219877, ALK: QT00028847, GAPDH: QT00079247).

### Reagents

#### Antibodies

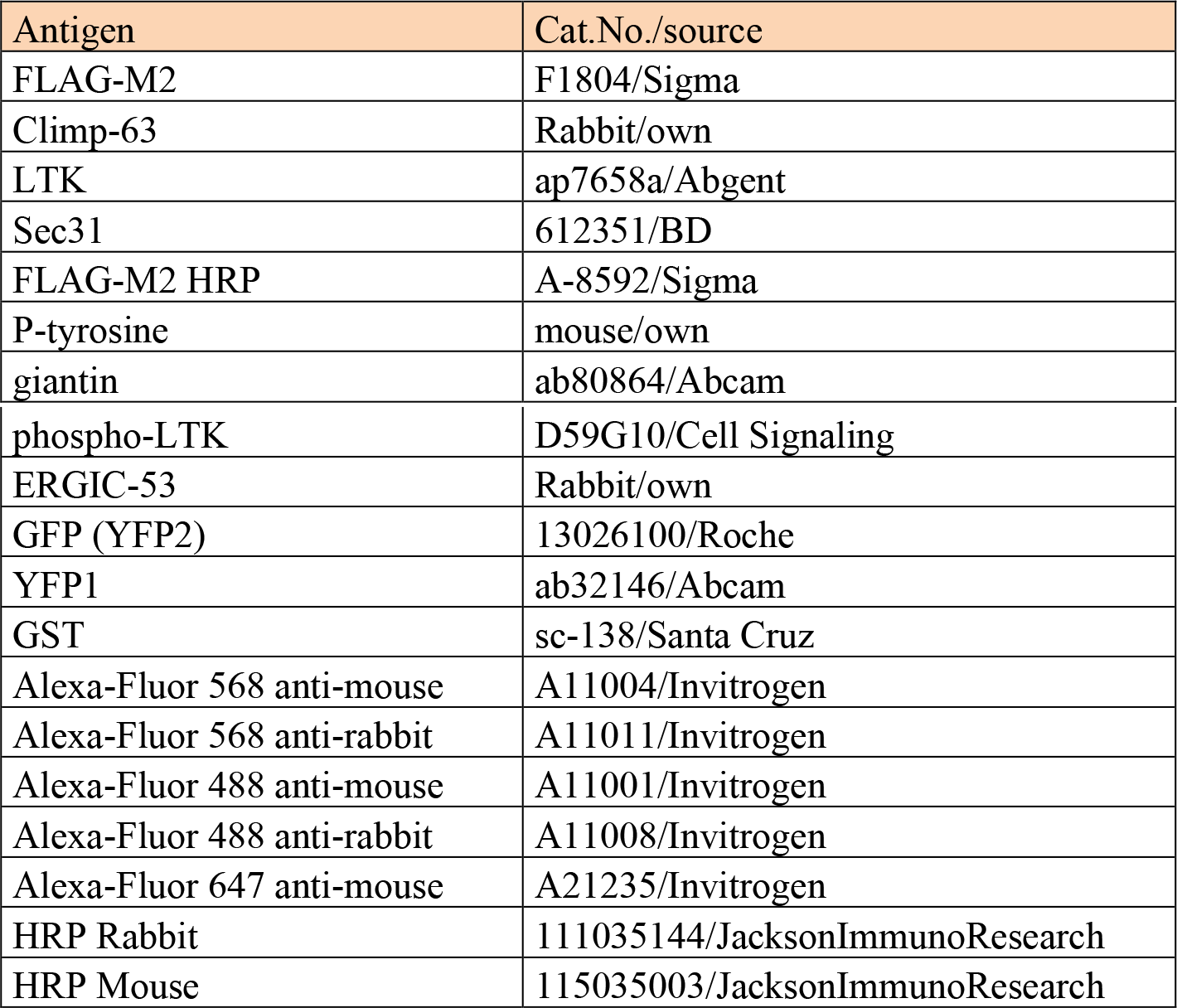

#### Primers

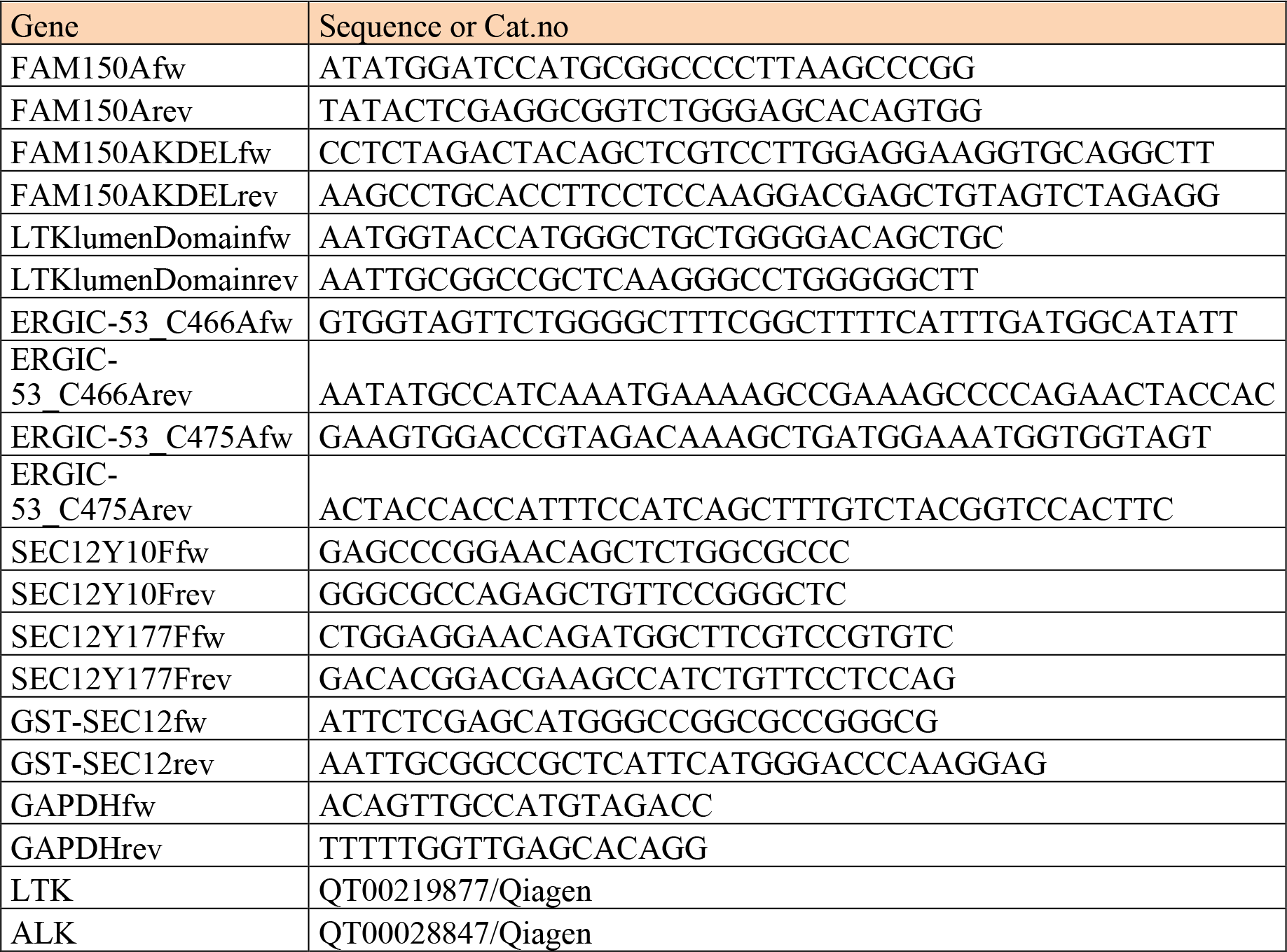

## Acknowledgement

Work in the Farhan lab was supported by grants from the Norwegian Research Council (NFR), the Norwegian Cancer Society (Kreftforeningen), the Anders Jahre Foundation, the Rakel-Otto-Bruun Legat, the Swiss Science Foundation (SNF) and the German Science Foundation (DFG). FGC is supported by a PhD scholarship from the Institute of Basic Medical Sciences, University of Oslo. CB was funded by the Boehringer Ingelheim Foundation.

